# Flight vocalizations and displays of Chimney Swifts (*Chaetura pelagica*): description and possible functions

**DOI:** 10.1101/2025.08.16.670661

**Authors:** Louise A. Peppe, Gary Ritchison

## Abstract

The vocalizations of many songbirds have been well documented and analyzed, but less is known about the vocal behavior of many non-passerines, including swifts. When flying alone and during aerial displays with conspecifics, Chimney Swifts (*Chaetura pelagic*) often utter a twitter call consisting of a series of high-frequency chip notes. However, little is known about the possible function(s) of swift flight displays and their twitter call. Our objectives were to record, analyze, and document the aerial behaviors and associated vocalizations of adult Chimney Swifts. We studied in Madison County, Kentucky, where they used abandoned, concrete shelters for roosting and nesting. Camcorders were used to record swift behavior and vocalizations during the 2008 and 2009 breeding seasons (April – September). We examined possible variation in the characteristics of swift vocalizations and the frequency of different aerial behaviors among breeding stages and behavioral contexts. Chimney Swifts engaged in more interactive pair flights during the nest-building/egg-laying stage, when females are likely fertile, and significantly more than during the pre-building and nestling stages, suggesting the possible importance of pair flights in courtship, pair bonding, and mate-guarding. Our results also suggest that V-ing (a raised-wing display) may be important in establishing or maintaining pair bonds because swifts engaged in this behavior more frequently during close chases involving two birds. We separated the typical swift twitter into two bouts: “steady” bouts and “quick” bouts. Mean chip rates were higher for the quick portion of the call, but we found no differences in the use of steady and quick bouts among nesting stages or in different behavioral contexts. Mean chip rates for quick bouts were highest for single birds and lowest for two and three birds, suggesting that twitter calls provide information about a bird’s location; to help coordinate movements while flying near others (e.g. when foraging and during chases), We were unable to identify individual swifts; such identification would facilitate the investigation of individual variation in call characteristics (e.g. chip rate and steady/quick bout rate) and relationships between and among swifts engaging in different activities and flight displays.

---

Little is known about the vocal behavior of swifts (Apodidae) because they typically spend much of the day in wide-ranging, rapid flight (Cink and Collins 2002), and often nest and roost in difficult-to-access locations. Vocalizations uttered by swifts in flight have been recorded and described, but most studies of swift vocalizations have been observational and the possible function(s) of those vocalizations is understudied. However, investigators have often documented the contexts in which calls were uttered and inferred functional significance. For example, Black Swifts (*Cypseloides niger*) sometimes emit a low-pitched call consisting of a rapid series of low, flat, twittering chips (Sibley 2000) as well as a sharp *cheet* note when adults approach their nest sites (Gunn et al. 2023). Marín (1997) observed that Black Swifts also uttered soft high-pitched sounds during group chases and high-pitched rolling twitters during pair chases and high speed dives involving pairs, and suggested that group and pair chases may play a role in pair formation and pair bond reinforcement.

The vocal repertoire of White-throated Swifts (*Aeronautes saxatalis*) includes a long, drawn-out rattling or twittering call, a loud, sharp, single note call, and a 2-note flight call, but little is known about the possible functions of these calls (Ryan and Collins 2003). Vaux’s Swifts (*Chaetura vauxi*) utter high-pitched, rapid chipping and buzzy insect-like twitters as well as high-pitched squeals or squeal-like sounds, but the function of these calls remain to be determined (Schwitters et al. 2021).

Common Swifts (*Apus apus*) have a long screaming call, a duet screaming call, and a nest call (Bretagnolle 1993, Oudheusden 2006). The long screaming call, presumably uttered by males, may have a territorial or agnostic function (Bretagnolle 1993), and duetting may be important for pair bonding and territorial defense (Malacarne et al. 1991). Nest calls may play a role in establishing pair bonds (Bretagnolle 1993). Pallid Swifts (*Apus pallidus*) also engage in duets (Malacarne and Cucco 1990, Cucco et al. 1994) that may aid in reproductive synchronization and competition among males for females (Cucco et al. 1994).

As with other species of swifts, little is known about the vocal behavior of Chimney Swifts (*Chaetura pelagica*). When flying, these swifts utter buzzy insect-like rolling twitter calls (also called chipper calls; Steeves et al. 2020) that consist of a series of high-pitched chip notes. The interval between chip notes reportedly varies (Bent 1940, Steeves et al. 2020), but the possible functional significance of such variation is unknown. During the non-breeding season, Bouchard (2005) observed that Chimney Swifts vocalize most when entering and exiting roosts in the fall and suggested that they track each other’s positions with non-overlapping chip notes. Overlapping calls may help swifts synchronize movements and maintain group cohesion. Bent (1940) and Fischer (1958) suggested that vocalizations uttered during aerial displays may play some role in courtship and the formation of pair bonds.

No one to date has examined the characteristics of Chimney Swift calls uttered at different times during the breeding season and in different behavioral contexts. Such information could provide insight concerning the possible functions, as well as the possible function of variation in the characteristics (e.g., variation in inter-note intervals), of these calls. Thus, our objectives were to record, analyze, document, and quantify the aerial behaviors and associated vocalizations of adult Chimney Swifts and to examine how vocalizations and aerial behaviors vary among breeding stages and in different behavioral contexts.

## Methods

Field work took place from late April to mid-September of 2008 and 2009 on the Blue Grass Army Depot (BGAD), Madison County, Kentucky. Chimney Swifts roosted and nested in small concrete shelters (*N* = 54; 4.6 m x 2.4 m, with two 0.9 m x 2.4 m entrances) on the BGAD. The mean distance between nearest shelters was 447 ± 194 m. Field observations began in mid-April when swifts arrived at the BGAD and began pair formation (Steeves et al. 2014). Swifts were observed at 27 shelters during 2008 and 25 shelters during 2009. Nests were never built in three shelters during 2008 and one shelter in 2009 so observations of swifts at these shelters were not included in our analyses. To monitor nest status, we visited shelters at intervals of 1-10 days and recorded presence or absence of a nest and the number of eggs and nestlings present. We used these data to categorize nest stages as pre-building, building, building-laying, incubating, nestling, and post-fledging. The pre-building stage began when swifts were first observed flying near a shelter, but nest construction had not yet started. Building began as soon as twigs were found attached to a shelter wall. The laying stage began when the first egg was observed in a nest.

However, because swifts continue to add twigs to nests well into the egg-laying stage, this stage was called the building-laying stage. The incubation stage began with clutch completion and the nestling stage began when at least one hatchling was present. The post-fledging stage began when all swifts had left the nest shelter.

The behavior and vocalizations of Chimney Swifts were recorded on videotape using a camcorder (Handycam, Sony, Tokyo, Japan) with an internal microphone. We video-recorded swifts an average of 2.9 ± 2.1 (SD) times per shelter from 22 May 2008 to 6 September 2008 and 2.0 ± 1.5 times per shelter from 15 May 2009 to 2 August 2009. During periods with high ambient noise in 2008 (when many periodical cicadas [*Magicada* spp.] emerged), a directional microphone (ME-66, Sennheiser) was attached to the camcorder to improve the quality of recordings; an external microphone was not needed in 2009. We video-recorded swifts at different times during the day, excluding the period from 13:00 to 15:00 when the sun was directly overhead and observing swifts was difficult.

The field of view on videos was limited to an area around swifts that varied with distance and the extent to which we varied camcorder settings while video-recording (i.e., between narrow and wide fields) so, when viewing videos, we could not determine if any swifts were present outside the camcorder’s field of view (portion of sky captured on videotape). Swifts entering our field of view were usually observed for about 2 to 20 sec. If multiple swifts were present in an area, we attempted to videotape all individuals (e.g., pairs or groups) flying together rather than focusing (i.e., narrowing the field of the camcorder) on birds flying alone. When birds were engaged in chasing behavior near a shelter, we assumed that at least one of the birds involved was a member of the resident pair nearest the shelter where we were videotaping. However, given how fast and how high swifts typically fly, we were not able to confirm the identity of swifts in the video-recordings.

### Analysis of aerial behavior

Based on a review of videotapes, we categorized Chimney Swift aerial behavior as either (1) non-interactive or (2) interactive. Non-interactive behaviors included a single swift flying alone, a single swift flying into a shelter, and group flying. A swift flying alone was likely foraging with no other swifts consistently present in our field of view or the field of view of the video-recorder. During group flying, two or more swifts were observed in flight within our field of view, likely foraging, but they did not appear to be following or interacting with each other.

Swifts exhibited the following interactive aerial behaviors, all of which, excluding tumbling falls, have been described in detail by Fischer (1958): (1) group chases, (2) pair chases, (3) trio chases, (4) V-ing, (5) tumbling falls, and (6) apparent contact. In addition to noting these behaviors and associated vocalizations, we sometimes video-recorded one or more swifts flying into the shelters where their nests were located.

Chasing behavior (group, trio, and pair) included all interactive behaviors and synchronized-flight events. First described by Bendire (1895), this behavior has since been explained and categorized in more detail. Group chases involve at least four birds, and were called a “loose association” by Fischer (1958). During a group chase, three or more swifts closely follow one leading bird in a high-speed, synchronized flight. Occasionally, one swift might leave the group or another might join the group. As the swifts separate, pair or trio chases often begin. In some cases, a pair or trio leave a group and initiate a chase. Trio chases, first documented by Kingston (1891) and MacNamara (1918), are characterized by two swifts following one leading bird. When most intense, these flights involved the most rapid flight, greatest heights, longest distances of any swift aerial displays, and can sometimes last for more than 5 min (Fischer 1958). During trio chases, the distance between the first two birds was often about twice the distance between the second and third birds (pers. observ.).

However, as flight speed increases, distances between swifts became more similar (Fischer 1958, pers. observ.). Pair chases, referred to as “flying together” or “flying in association” by Fischer (1958), were high-speed, synchronized flights involving two birds.

All interactive chases (group chases, trio chases, and pair chases) were further categorized as either close (0-5 m between swifts) or distant (>5 m between swifts) based on the distance between swifts. Approximate distances between swifts were determined when reviewing video-recordings. We used the length of swifts (∼12 cm; Cink and Collins 2002) on the screen as a guide for estimating the approximate distance between swifts.

Close chases were pair chases, trio chases, or group chases, with swifts often separated by <1 m, but never by > 5 m for more than a few seconds (e.g., as they flew through the branches of trees). Close chases often involved V-ing (see below). During distant chases, swifts were usually about 10 to 20 m apart, but never < 5 m.

During chases, one or two swifts sometimes engaged in V-ing (Fischer, 1958). During these displays, one bird suddenly raised its wings to form a sharp angular “V” shape. The other bird sometimes, but not always, responded by V-ing as well. Fischer (1958) stated that the trailing bird always initiated this display, but we observed many instances when the leading bird was the first to raise its wings. Both birds then glided together in a downward dive. V-ing displays never involved more than two birds (Fischer 1958), were most common during pair chases, and occasionally occurred during a trio chase.

Tumbling falls occurred when two swifts came together, locked their feet, and began tumbling toward the ground. This behavior has been documented only twice previously in Chimney Swifts (Cink and Collins 2002) and was observed once during our study. The function of this interaction, which occurs more frequently in other swift species (Bradbury 1918, Dawson 1923, Michael 1926, Bent 1940, Marin 1997, Ryan and Collins 2003), is unclear (Marín and Stiles 1992).

Infrequently, Chimney Swifts may engage in aerial copulations (Fischer 1958). This occurs as two swifts come together when V-ing, with the male slightly above and behind the female. The male makes momentary contact, thrusting his body forward without moving the wings. Although there have been a few reports of aerial copulations (Bagg and Elliot 1937, Bent 1940), it is not clear if they actually occur because swifts may appear to come in contact during V-ing displays (Fischer 1958, pers. observ.). There are reported cases of aerial copulation by Common Swifts (*Apus apus*; Chantler and Driessens 1995), and cases of apparent aerial copulation in Vaux’s Swifts (Schwitters et al. 2021), but Chimney Swifts more commonly copulate at nest sites (Steeves et al. 2020). During our study, one V-ing bird occasionally flew close enough to another (often V-ing) bird during pair chases so that they appeared to make contact. However, because we could not be certain that the swifts actually made contact during these interactions, we referred to this behavior as apparent contact.

We obtained 795 minutes of usable video-recordings of swift aerial behaviors and vocalizations during 2008 and 2009. All video-recordings were subsequently reviewed and, for each aerial behavior, we noted the total time swifts engaged in the behavior, the date, shelter number, and breeding status of swifts nesting in the nearest shelter.

### Analysis of vocalizations

All video recordings were digitized and the corresponding sound files were reviewed and analyzed using Raven Interactive Sound Analysis Software (Cornell Lab of Ornithology 2011). Sound clips contained periods of silence, solitary chip notes, and periods of consistent calling that we termed calling bouts. A solitary chip was defined as any chip that was not part of a calling bout, typically separated from other chips by >1.5 sec. Calling bouts, previously categorized as uniform calls and varying calls (Bouchard 2005), were classified as steady or quick. Steady calling bouts were defined as bouts with relatively consistent and predictable inter-note durations (duration between chip notes), whereas quick calling bouts were bouts with less consistent inter-note durations (Figure 1). Inter-note durations for steady bouts were longer than those for quick bouts. Calling bouts containing both steady and quick segments of calling were categorized as two different bouts (e.g., Figure 1).

**Figure 1.**
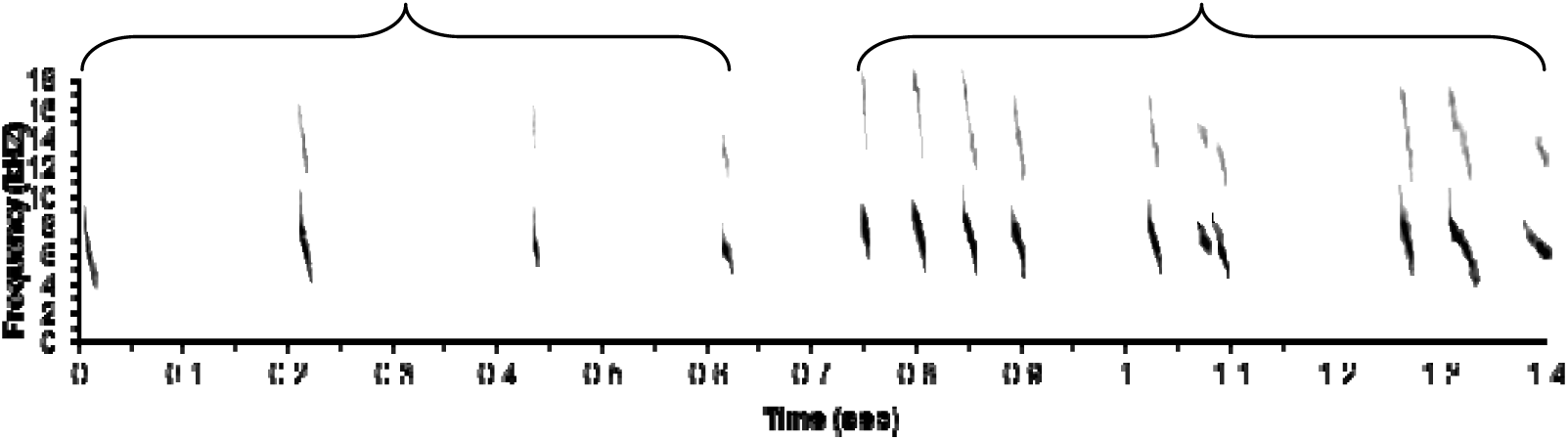
Examples of a steady calling bout and a quick calling bout. The entire clip is representative of a typical Chimney Swift twitter.

All recorded swift vocalizations were analyzed relative to their aerial behaviors. For every aerial behavior, we recorded the duration of each sound clip, number of calling bouts and solitary chip notes, the date, shelter number, and breeding status of swifts at the nearest shelter. A subsample of calling bouts was analyzed in more detail. We used Microsoft Excel’s (2004) random-number generator to select the calling bouts to be analyzed. For analysis of calling bouts, non-interactive aerial behaviors of flying alone and group flying were grouped together as non-interactive flight because vocalizing swifts may have been present outside the camcorder’s field of view.

For every sampled calling bout, we recorded the duration (in seconds) of the bout, number of swifts calling, number of chips, chip rate, and type of calling bout. Date, shelter number, and breeding status were also noted for each bout. The number of birds was determined by looking for distinguishing characteristics in the chips given by individual birds including differences in the amplitude, frequency, or morphology of chip notes.

### Statistical analyses

We used repeated measures analysis of variance (ANOVA; SAS Institute 2014) to determine if breeding stage (pre-building, building, building-laying, incubating, nestling, and post-fledging) had a significant effect on the percent of time swifts engaged in interactive and non-interactive aerial behaviors. The repeated measures procedure was used because we observed birds at each shelter multiple times. When necessary, Tukey’s post-hoc tests were used to determine differences among means of different breeding stages.

Repeated measures analysis of variance was also used to determine if swifts engaged in any type of interactive behavior (close-group chases, distant-group chases, close-trio chases, distant-trio chases, close-pair chases, distant-pair chases, V-ing, and apparent contact) significantly more than any other interactive events. For analysis of these data, the number of times swifts engaged in each behavior was converted into rate, i.e., the number of events per minute.

We also examined possible differences in the frequency of use of quick versus steady bouts during different nesting stages and during different detailed behavioral contexts (flying alone, close-group chase, distant group chase, close-pair chase, distant-pair chase, close-trio chase, and distant-trio chase). For this analysis, we determined the number of quick and steady bouts during each nest stage or interactive behavior at each shelter and, for all cases where swifts uttered at least three bouts per nest stage or interactive behavior, we determined the percentage of total bouts that were quick or steady bouts. After transforming the percentage data to generate a normal distribution (arcsine square root), we used repeated measures analysis of variance to examine possible differences among nest stages and interactive behaviors in use of quick versus steady bouts.

Repeated measures ANOVA was used to determine if the number of birds (one, two, or three) chipping affected chip rates. Tukey’s *post-hoc* tests were used to determine how the number of birds chipping influenced chip rates. Repeated measures ANOVA was also used to determine if chip rate differed by type of bout (steady or quick) or nest stage, and if behavioral context (interactive or non-interactive), detailed behavioral context, or grouped behavioral context (flying alone, group chase, pair chase, and trio chase) influenced chip rates.

For analysis of aerial behavior and vocalizations, we assumed that repeated measurements of swifts at a particular shelter were not independent. Means are reported ± SE, and results of P ≤ 0.05 were considered statistically significant.

## Results

We analyzed 212 non-interactive flight bouts. These included 41 shelter entrances along with instances of swifts flying alone and in groups. Numbers of each were not recorded due to the nature of non-interactive flight (swifts were often not flying close enough together to capture in the camera’s field of view); therefore, it was unknown whether other swifts were present outside the camera’s field of view. In addition, we analyzed 518 interactive flight bouts, including 72 close-group chase bouts, 24 distant-group chase bouts, 152 close-pair chase bouts, 96 distant-pair chase bouts, 129 close-trio chase bouts, and 45 distant-trio chase bouts. The number of bouts sampled per behavior was indicative of the total number of bouts of each aerial behavior, e.g., we observed fewer calling bouts during distant-group and distant-trio flights.

### Aerial behavior and breeding stage

The percent time swifts engaged in non-interactive flights when alone (*F*_5,33_ = 1.2, *P* = 0.32) and participating in group flights (F_5,33_ = 1.3, P = 0.29) did not differ among breeding stages. Similarly, for interactive flights, the percent of time swifts engaged in trio flights (*F*_5,33_ = 0.7, *P* = 0.60) did not differ among breeding stages. However, the percent of time swifts were engaged in interactive group flights (close and distant combined; *F*_5,33_ = 2.7, *P* = 0.004) and interactive pair flights (close and distant combined; *F*_5,33_ = 2.7, *P* = 0.036) did vary with breeding stage. Swifts spent more time engaged in interactive group flights during the post-fledging stage (Figure 2), and more time engaged in interactive pair flights during the nest building/laying stage than during the prebuilding and nestling stages (Figure 3).

**Figure 2.**
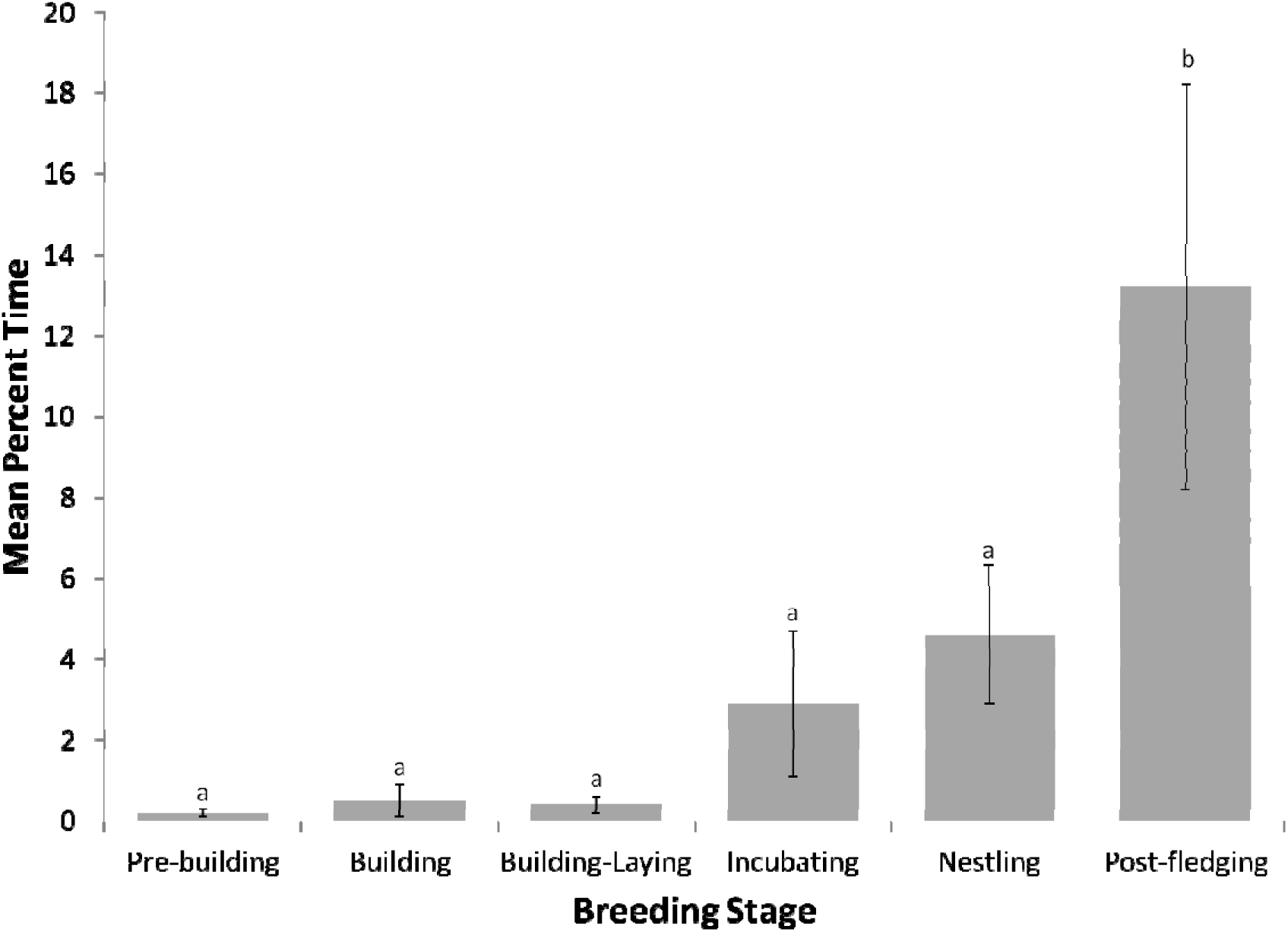
Mean percent time (± SE) Chimney Swifts engaged in interactive group flights during each breeding stage. Means with the same letter were not significantly different.

**Figure 3.**
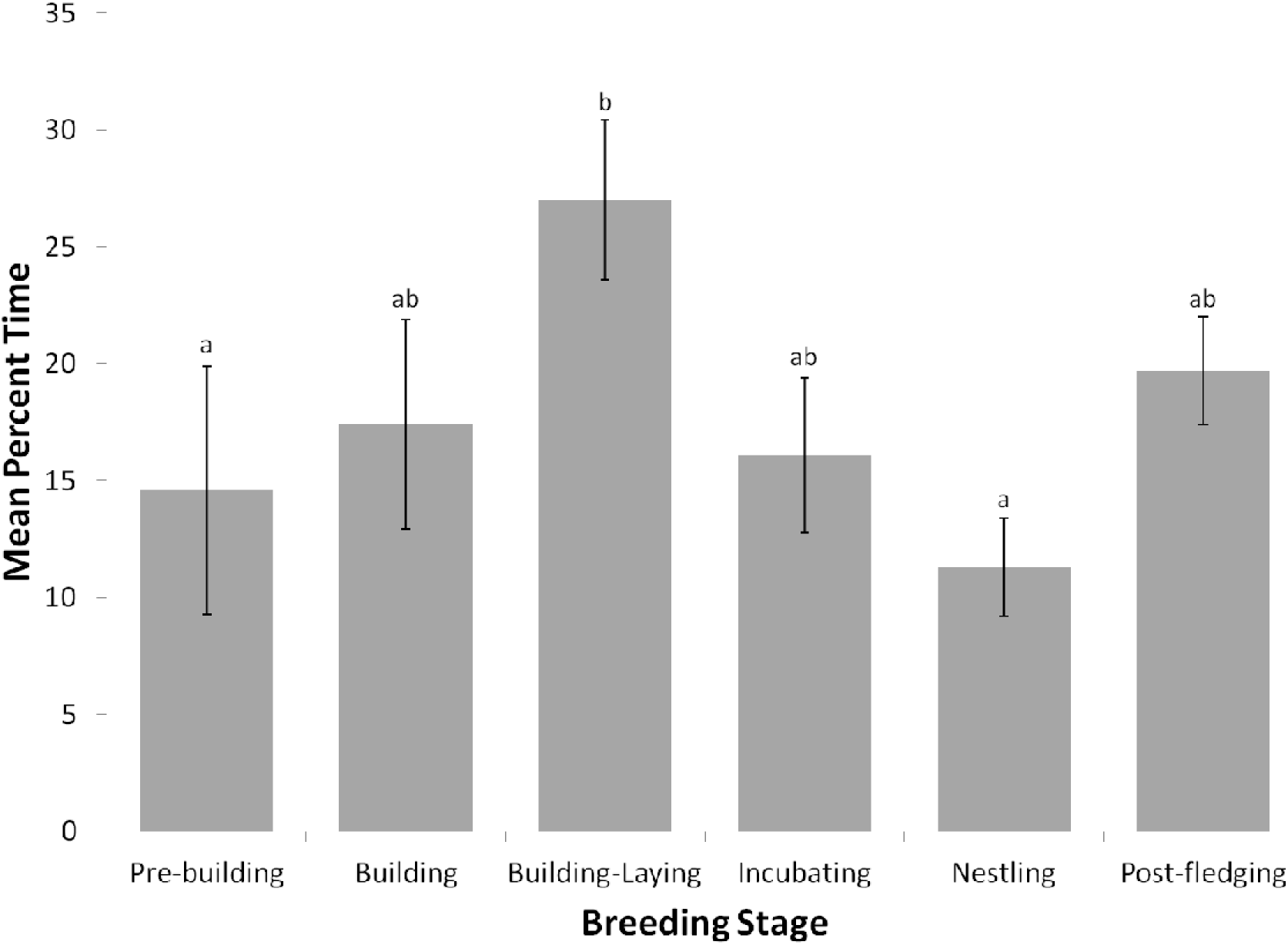
Mean percent time (± SE) Chimney Swifts engaged in interactive pair flights during each breeding stage. Means with the same letter were not significantly different.

We found no significant differences among breeding stages in the rates at which swifts engaged in any specific interactive behaviors, including close-group chases (*F*_5,33_ = 1.3, *P* = 0.30), distant-group chases (*F*_5,33_ = 0.6, *P* = 0.68), close-trio chases (*F*_5,33_ = 0.7, *P* = 0.61), distant-trio chases (*F*_5,33_ = 0.4, *P* = 0.84), close-pair chases (*F*_5,33_ = 1.3, *P* = 0.30), distant-pair chases (*F*_5,33_ = 1.2, *P* = 0.32), and V-ing (*F*_5,33_ = 1.6, *P* = 0.19). Although not significant, most observations of V-ing were during close-pair chases (80 of 101, or 79.2%), with 10 observations of V-ing during close-group chases (*F*_5,33_ = 1.6, *P* = 0.19) and 11 during close-trio chases. Similarly, most observations of apparent contact occurred during close-pair chases (10 of 11, or 90.9%); one observation of apparent contact occurred during a close-group chase. In addition, most observations of apparent contact were made during female fertile periods (8 of 11, or 72.7%); other observation of apparent contact were made during female post-fertile (*N* = 2) and pre-fertile (*N* = 1) periods.

### Vocalizations – chip and bout rates

When one swift was vocalizing, mean chip rates of quick bouts (*N* = 279, 14.2 ± 0.2 chips/sec) were higher (*F*_1,24_ = 650.0, *P* < 0.0001) than mean chip rates of steady bouts (*N* = 232, 5.0 ± 0.1 chips/sec). Similarly, when two or more birds were vocalizing, mean chip rate of quick bouts (*N* = 104, 10.4 ± 0.2 chips/sec/bird) was higher (*F*_1,13_ = 221.0, *P* < 0.0001) than that of steady bouts (*N* = 74, 4.8 ± 0.1 chips/sec/bird).

For steady bouts, mean chip rates of single birds (chips/sec) and multiple (i.e., 2 or 3) birds (i.e., chips/sec/bird) did not differ (*F*_1,15_ = 0.7, *P* = 0.41). However, for quick bouts, mean chip rates of single and multiple birds did differ significantly (*F*_1,16_ = 82.6, *P* < 0.0001), with a mean rate of 14.2 ± 0.2 chips/sec (*N* = 278) for single birds and 10.4 ± 0.2 chips/sec/bird for two or three birds.

For swifts flying alone, chip rates for steady bouts did not vary among nesting stages (*F*_5,27_ = 1.5, *P* = 0.23), but, for quick bouts, chip rates did vary with nest stage (*F*_5,30_ = 3.9, *P* = 0.008). For quick bouts, chip rates of single birds were significantly lower (mean = 12.1 ± 0.7 chips/sec, *N* = 23) during the prebuilding period than during the building (mean = 14.9 ± 0.4 chips/sec, *N* = 48), building/laying (mean = 14.9 ± 0.3 chips/sec, *N* = 79), incubation (mean = 14.5 ± 0.4 chips/sec, *N* = 45), nestling (mean = 13.8 ± 0.3 chips/sec, *N* = 59), or post-fledging (mean = 13.2 ± 0.5 chips/sec) periods (Tukey’s test, P < 0.05). For multiple swifts (2 or 3), chip rates did not differ among nesting stages for either steady bouts (*F*_5,5_ = 1.5, *P* = 0.33) or quick bouts (*F*_5,9_ = 0.8, *P* = 0.58).

For both quick and steady bouts, chip rates did not differ with context (interactive vs. non-interactive) for either single birds (steady: *F*_1,14_ = 1.0, *P* = 0.33; quick: *F*_1,19_ = 0.02, *P* = 0.88) or 2 or 3 birds (steady: *F*_1,5_ = 2.0, *P* = 0.21; quick: *F*_1,6_ = 0.9, *P* = 0.38). Similarly, chip rates did not differ among detailed behavioral contexts (flying alone, close-group chase, distant-group chase, close-pair chase, distant-pair chase, close-trio chase, and distant-trio chase) for either steady bouts (*F*_6,57_ = 1.1, *P* = 0.40) or quick bouts (*F*_6,67_ = 0.9, *P* = 0.51). Finally, chip rates did not differ among grouped behavioral contexts (flying alone, group chase, pair chase, and trio chase) for either steady (*F*_3,36_ = 1.1, *P* = 0.35) or quick (*F*_3,43_ = 1.1, *P* = 0.37) bouts.

We found no difference among nesting stages in use of quick versus steady bouts (*F*_5,21_ = 2.1, *P* = 0.11). Among detailed behavioral contexts, analysis revealed a significant difference in use quick versus steady bouts (*F*_6,47_ = 2.5, *P* = 0.035; Figure 4). However, post-hoc analysis revealed a single significant difference between behavioral contexts, with swifts using more quick bouts during close-group chases than during distant-trio chases (Tukey’s test, *P* < 0.05).

**Figure 4.**
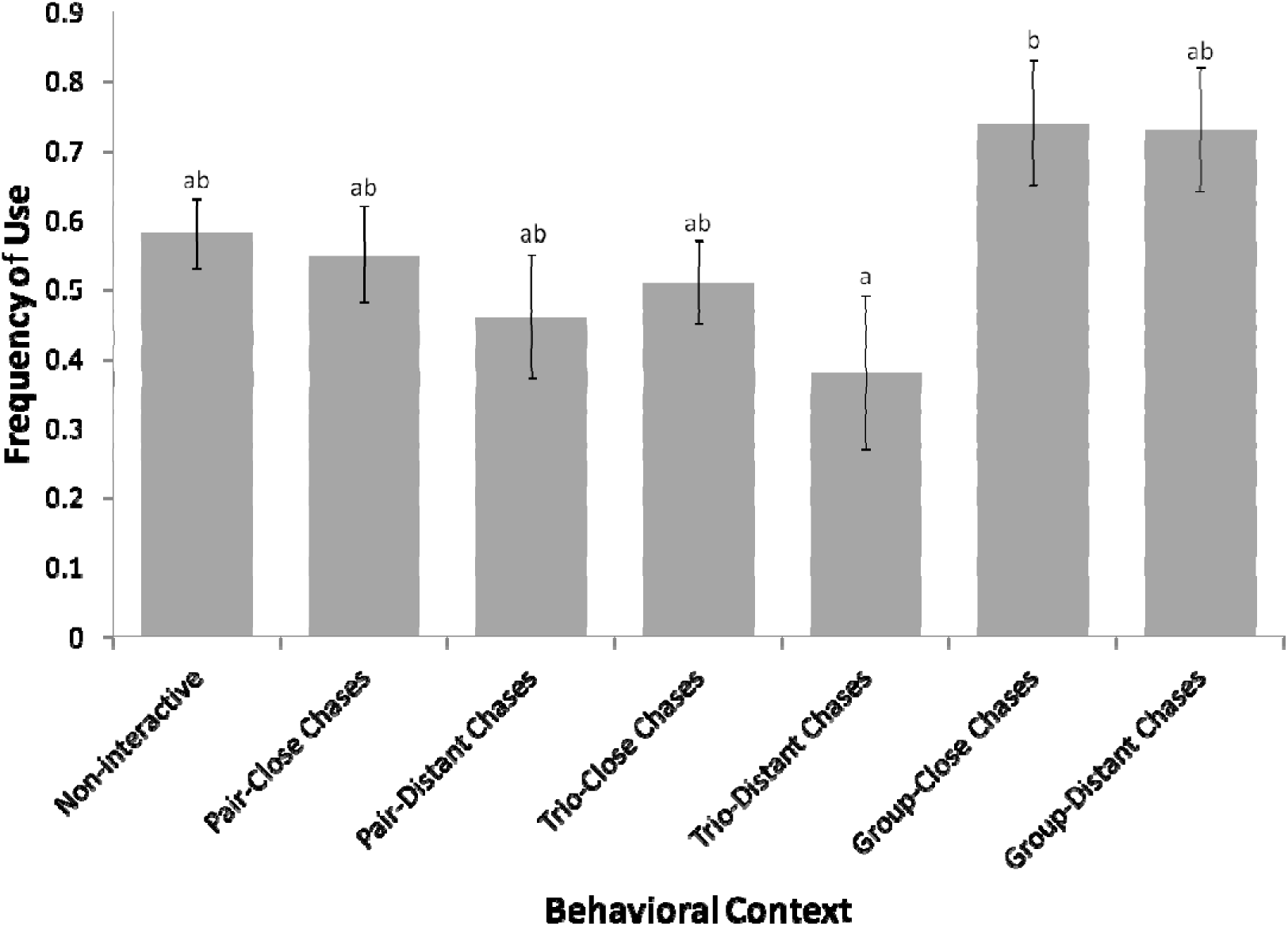
Mean frequency (± SE) of use of quick bouts relative to steady bouts by Chimney Swifts. Means with the same letter are not significantly different.

**Figure 5.**
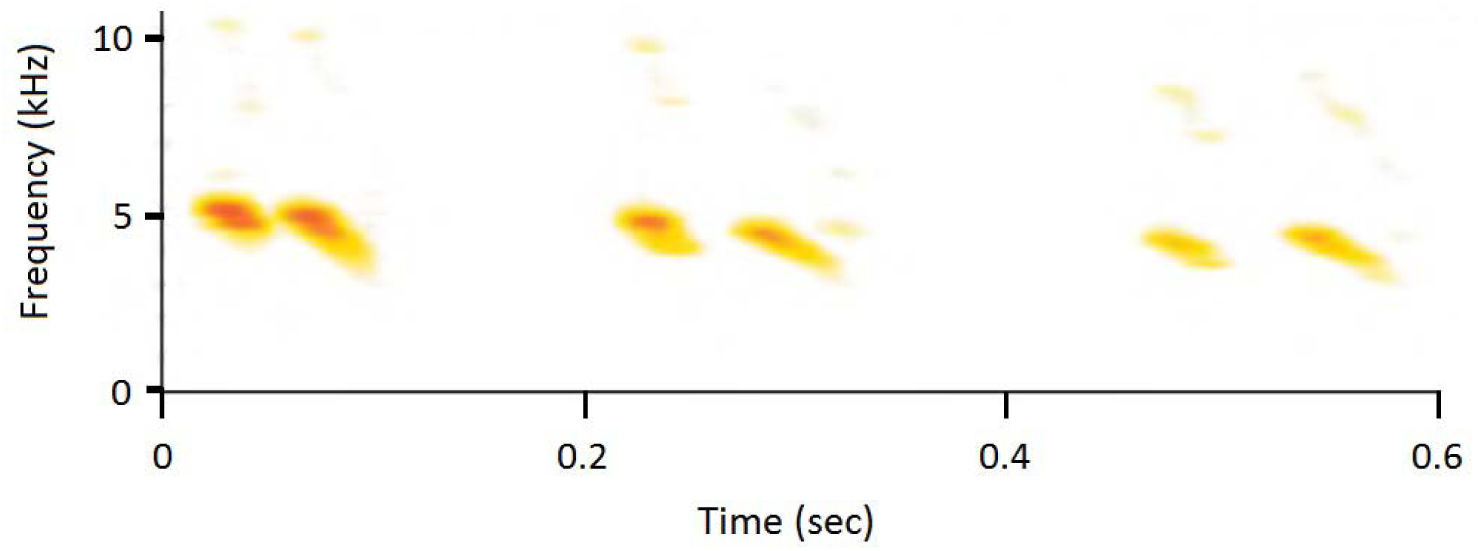
Sonagram of the chip notes of two Chimney Swifts flying near each other. Note that the notes uttered by the two swifts do not overlap.

## Discussion

### Aerial behavior and breeding stage

We found that the percent time spent by Chimney Swifts in non-interactive flights either when alone or in groups did not vary among breeding stages. During these non-interactive flights, Chimney Swifts were likely foraging. Unfortunately, we were not able to identify individuals in flight so no conclusions can be drawn concerning how time spent foraging by individual swifts might vary with breeding stage. However, Zammuto et al. (1981) found that, during the nestling stage, adult Chimney Swifts provisioned nestlings about every 15 minutes, and that the mean interval between visits to nests by adults varied significantly with nestling age and brood size. Such results suggest that time spent foraging by adult Chimney Swifts, as opposed to time spent flying, but not actively foraging, likely does vary with breeding stage.

Chimney Swifts spent more time engaged in interactive pair chases during the building-laying stage than during other breeding stages, and significantly more than during the pre-building and nestling stages. During the building-laying stage, pair bonding and synchronization is important and, in addition, females are likely fertile during this stage. In White-throated (Ryan and Collins 2003), Black (Marín 1997), and Vaux’s (Rathbun and Bent 1940, Schwitters et al. 2021) swifts, chases peak during courtship in the spring and early in the breeding season, suggesting an association with establishing and reinforcing pair bonds. For male Chimney Swifts, pair chases might also serve a mate-guarding function. The extent to which female Chimney Swifts might engage in extrapair copulations in unknown (Steeves et al. 2020). However, Martins et al. (2002) reported extra-pair young in four of 42 (9.5%)

Common Swift nests, suggesting that possibility of extra-pair behavior in other species of swifts (Apodidae). The increased frequency of pair chases by Chimney Swifts during the building-laying stage in our study suggests that, for males, one possible function of such behavior is mate guarding.

We also found that Chimney Swifts spent more time in group chases during the post-fledging period. Large groups of swifts often flew together at this point in the breeding season, possibly due to the presence of newly fledged young (pers. observ.). Among some species of swifts, including Chimney Swifts (Fischer 1958), adults and fledglings may return to their nest sites to roost (Bull and Blumton 1997), so families of swifts may also fly together during the day. In addition, swifts may begin roosting communally after the breeding season (Bull and Blumton 1997, Rioux et al. 2010) and birds in those roosts may also forage and fly in groups during the day.

We found no differences among breeding stages in the frequency of close and distant pair, trio, or group flights or chases. One possible function of these aerial interactions during the nestling period is that they could provide swifts with information about the location of insect prey. The abundance of aerial insects varies in space and time. For example, Brown (1985) noted that aerial insect prey of Cliff Swallows (*Hirundo pyrrhonota*) were ephemeral and could be concentrated by localized convection currents, the location of mating swarms, or mass emergences.

Swallows breed in colonies that can serve as information centers, i.e., unsuccessful foragers can observe birds returning to colonies to feed young, identify successful foragers based on the number of provisioning visits or the extent to which the bolus of captured insects cause throats to bulge, then follow them when they leave colonies to forage (Brown 1988). Swifts are not colonial, but when foraging for nestlings they do form captured insects into a bolus that can cause their throats to bulge (e.g., Lack 1956, Fischer 1958, Bull and Beckwith 1993). As such, by flying near others and noting the relative sizes of food boluses, swifts may be able to identify successful foragers that could be followed to good foraging locations. Prior to nesting, interactive flights may provide swifts with information about the mated status or condition of other swifts, including potential mates.

Due to small sample sizes, we were unable to statistically examine one type of aerial interaction, the V-ing display, but V-ing occurred most often (80 of 101) when swifts engaged in close-pair chases. This display, sometimes also referred to as the raised-wing display, has also been reported in other species of swifts, including Common Swifts (Lack 1956, Cramp 1985), Vaux’s Swifts (Schwitters et al. 2021), and White-throated Swifts (Ryan and Collins 2003). In Common Swifts, investigators have suggested that the raised-wing display is a precopulatory solicitation display (Lack 1956, Cramp 1985). Because Chimney Swifts copulate near nest sites rather than in flight, Fischer (1958) suggested that the V-ing display was important for physiological synchronization and maintenance of the pair bond. In support of this hypothesis, most V-ing displays in our study were observed during close-pair flights.

### Vocalizations – chip and bout rates

Chimney Swifts in our study uttered chip notes at significantly different rates during quick bouts and steady bouts. However, use of these two types of bouts did not differ among nesting stages and, in addition, we found minimal difference, with no apparent biological significance, in use of the two bout types among detailed behavioral contexts. Such results suggest that quick and steady bouts may be functionally equivalent. Bouchard (2005) suggested that one possible function of twitter calls is to provide information about a Chimney Swift’s location. While foraging and, especially during pair, trio, and group flights and chases, swifts fly at high speeds and twitter calls may be used to help coordinate movements, providing information that could be important for maintaining each bird’s position relative to that of others. In support of this hypothesis, we found that the rates at which chip notes were uttered during quick bouts when two swifts were flying near each other and calling were slower than the rates at which chip notes were uttered by swifts flying alone, suggesting that swifts deliberately slow down their call rates when flying near others. One possible explanation for this is that calling at a slower rate may reduce the likelihood of notes overlapping. Although not quantified, we found that, when two or more Chimney Swifts were flying together and more than one was calling, their chip notes often did not overlap (e.g., Figure 6). Non-overlapping notes would presumably make it easier for swifts to hear, and better judge the position of, nearby swifts.

For pairs of Chimney Swifts flying together, the coordination of calling rates to avoid note overlap is similar to the coordination that occurs in some species of birds when paired males and females sing duets. Investigators studying other species of swifts have suggested that coordinated calling by males and females may play an important role in breeding. For example, the duet screaming call of Common Swifts likely functions as a territorial call as well as in partner identification and synchronizing behavior (Bretagnolle 1993). Similarly, the ‘duetting calls’ of Pallid Swifts may be important for reproductive synchronization and pair-bond consolidation (Cucco et al. 1994) as well as territory defense (Malacarne and Cucco 1990).

Although Chimney Swifts in our study consistently uttered chip notes at different rates during quick and steady bouts, our analysis revealed no apparent difference in the respective functions of these two types of bouts. However, we were not able to identify individual swifts and could not determine the specific reason(s) why swifts were flying together. As a result, we were not able to examine the possibility that differences in social relationships between and among swifts (e.g., pair vs. non-paired swifts) or that swifts engaged in different activities (e.g., foraging vs. interacting for other reasons) might influence use of the two types of bouts in different contexts. Additional study will be needed to further examine use of the two types of bouts and determine possible differences in function.

For multiple swifts (2 or 3) flying together, we found no effect of nest stage on chip rate for either steady or quick bouts. For swifts flying alone, chip rates of steady bouts did not vary with nest stage, but, for quick bouts, chip rates were significantly lower (mean = 12.1 notes/sec) during the pre-nest-building period than during other nest stages (range of means = 13.2 – 14.9 notes/sec). The difference in mean chip rate between the pre-nest-building period and other nest stages ranged from just 1.1 notes/sec (pre-building stage vs. post-fledging stage) to 2.8 notes/sec (pre-building stages vs. both building and building/laying stages). The biological significance of such differences is unclear. However, the pre-building period is likely when pairs are forming and a slightly lower chip rate could be important if, for example, non-overlapping duets play a role in pair formation. If so, a slower chip rate during quick bouts by solitary swifts during the pre-building period could indicate availability to engage in non-overlapping duets, i.e., notes uttered at slower rates would be less likely to overlap those of another swift.

For both quick and steady bouts and for both swifts flying alone and with other swifts, chip rates did not differ with context (interactive vs. non-interactive), detailed behavioral contexts (non-interactive flight, close-group chase, distant-group chase, close-pair chase, distant-pair chase, close-trio chase, and distant-trio chase), or grouped behavioral contexts (non-interactive flight, group chase, pair chase, and trio chase). In contrast to our results, Bent (1940) and Steeves et al. (2020) suggested that the interval between the chip notes of Chimney Swifts can vary in different contexts, but, in neither case, did the authors provide data to support that suggestion. Thus, although our results suggest that chip rates do not vary with context, and only to a limited degree among nest stages, and, therefore, that Chimney Swifts do not vary those rates to convey different information in different contexts, as noted previously, additional study with marked birds is needed to better understand the possible role of chip rate in communication between known individuals and pairs.

In sum, although our results confirm the importance of the aerial displays of Chimney Swifts during the breeding season, we found little evidence that use of quick versus steady bouts or the rate at which notes are uttered during those bouts conveys specific information to conspecifics. Rather, our results suggest that Chimney Swifts may use their twitter calls primarily to help coordinate their movements during their high speed flights.

## Acknowledgments

We thank the U. S. Army for allowing us to conduct our study on their property.

Conflict of interest statement

The authors declare that they have no conflicts of interest.

## Funding

We thank the Department of Biological Sciences (Eastern Kentucky University) for financial support.

## Permits and ethics protocols

Permission to conduct our study was provided by the Eastern Kentucky University Institutional Animal Care and Use Committee.

## Statement on use of AI

Generative AI was not used in the production of this manuscript.

## Notes

### Competing Interest Statement

The authors have declared no competing interest.

